# Plant-Pollinator Interactions along the Altitudinal Gradient in *Berberis lycium* Royle: An Endangered Medicinal Plant Species of the Himalayan Region

**DOI:** 10.1101/2024.09.04.611343

**Authors:** Nahila Anjum, Sajid Khan, Susheel Verma, Kailash S. Gaira, Balwant Rawat, Mohd Hanief, Nakul Chettri

## Abstract

Mountain ecosystems influence species distribution by offering climatic variables intertwined with rising altitude. These climatic factors determine species phenology and niche width. Although the distributional patterns of some prominent insect groups in relation to altitude have been determined, the environmental preferences along the altitudinal range that differentially influence the pollination of specific plant species are unknown. Here we assess how the composition and abundance of pollinator fauna of the important medicinal plant *Berberis lycium* Royle (Berberidaceae) differ across five distinct altitudinal gradients (800-2200 m) in the Pir-Panjal mountain range in the northwestern part of the Indian Himalayas. We monitored insect pollinators of major groups (bees, butterflies, wasps, flies) over two consecutive flowering seasons during 2022-2023. In total, 39 insect species belonging to five orders and 17 families were observed visiting the plant species during the flowering period across the altitudinal gradient. The results of the linear regression model depict that all four pollination indices show a negative correlation with increasing altitude in foraging activities when all the data are pooled together. However, only foraging speed (FS) and index of visiting rate (IVR) were statistically significant. In the case of individual orders, only Lepidoptera exhibited a notable relation to altitude. However, asynchrony in foraging activities among other pollinator groups has been reported along altitudinal gradients. The reproductive output (fruit and seed production) shows a significant negative correlation with increasing altitude. We concluded that while altitude influences species distribution, it also differentially shapes plant-pollinator interactions, pollinator activities, and reproductive output. This work is of great significance in order to monitor plant-pollinator interactions, which are essential component of biodiversity rich but fragile mountain ecosystem.

## 1. Introduction

Mountain ecosystems are extremely fragile influenced by numerous drivers on species diversity and distribution patterns. They are home to about half of the world’s biodiversity hotspots and nurture numerous endemic species [1]. The Himalaya processes unique bio-climatic geographical ranges in Asia that separate the Tibetan plateau from the lowland plains of the Indian subcontinent. The degree of exposure to weather and environment varies along the different altitudinal zones [2]. Apart from the typical lapse rate in temperature conditions, there is substantial variation in wind speed and direction, orographic patterns, humidity, and precipitation, which differentially affect biodiversity at varying elevations [3].

For both plants and animals, including insects, patterns of species richness, and abundance often show a continually declining trend or a mid-elevation peak in response to elevation, however, these patterns differ between taxa and geographical regions [4-9]). These patterns of abundance and richness are predominantly shaped by abiotic variables physically bound to elevation [4, 10] that result in diversity-limiting factors at higher altitudes. On an average the Himalayas exhibit a standard temperature lapse rate of 5.65°C per 1000 m [11]. As a result, the higher altitude significantly alters the duration of the growth period [12] and metabolic rates [13] in insects. Unsuitable environmental conditions hamper their survival and altitudinal distributional range [14]. As the different potential pollinator insect groups appear to be impacted to varying degrees [15], communities of pollinators participating in the process of reproduction of mountainous angiospermic plants are likely to vary with elevation gradients [16].

The Pollinator assemblage of insect-pollinated plants remarkably exhibits temporal and spatial variations among plant communities [17] and sometime within a single plant species as well [18]. For example, orchids were pollinated by variable pollinators in different habitats and regions, for different preferences including nectar, pollen, trichomes, fragrance, lipids and resins [19] by a variety of morphologically distinct and specialized pollinators [20]. In addition, pollination is a vital regulating ecosystem service that underpins food production and contributes to gene flow and ecosystem restoration [21, 22]. Eventually the expected far-reaching impacts of the abiotic environmental factors on plant-pollinator interactions, we have little knowledge of how these interactions play out at altitudinal scales. Despite, the fact that several important plant species completely depend upon the insect pollinators for their survival, spatial variation in the pollinator groups along the altitudinal gradient is poorly understood.

Insects are the major pollinators of flowering plants in higher altitudes [23]. These insects are classified into several orders; the most common are Hymenoptera, Lepidoptera, Diptera, Hemiptera and Coleoptera [24]. Many researchers worldwide have studied the insect pollinator spectrum and pollination ecology of numerous medicinally important plants [25]. There are, however, few studies that assess all the necessary variables to ascertain how altitude influences the behaviours and range limits of species, including pollinator visiting rates, abundance, and reproductive output.. Altitudinal gradients provide the best opportunity for observing pollination ecology towards range limits as decline in pollination and resources may limit reproductive output at high altitudes [26, 27].

Berberidaceae is a diverse angiospermic family consisting of 17 genera and 650 species globally, most of which are located in the northern hemisphere. *Berberis lycium* Royle, an extremely useful shrub and most common member of the Berberidaceae family is native to the Himalayan region but may be found worldwide in subtropical and temperate climates. It is an extremely valuable medicinal plant that has been utilised in traditional medicine since time immemorial. Numerous medicinal uses of *B. lycium* were demonstrated, including wound healing, hepatoprotective, antioxidant, antibacterial, antihyperlipidemic and much more. In addition, the plant is traditionally used to treat ophthalmic, liver, skin, cough, and stomach problems. Because of its enormous nutritional and medicinal benefits, it is being overexploited for consumption and trade, which poses a serious threat to the existence of this plant in its natural habitat. The habitat of this plant is also being diminished by other anthropogenic activities like building and road construction, deforestation, agricultural road expansion in hilly areas, and so forth, which is causing a population decline [28, 29]. As per IUCN criteria of 2000, *B. lycium* has already been declared endangered in northwestern region of the Himalayas [30]. Therefore, conservation measures need to be taken to protect this species from additional threat in the near future.

In the northwestern Himalayas, *B. lycium* grows along a altitudinal gradient from 800 to 2200 m asl. Earlier, some studies [31] showed how climatic conditions could influence floral phenology and frequency, activity and life cycle of insect pollinators of a *Berberis* species. Owing to the wide range of habitat distribution of *B. lycium* and reproductive dependence of its insect pollinators [25, 30]. The present study is aimed to (1) document the diversity and variations of prominent insect pollinators of a common host plant species along altitudinal gradient; (2) compare the insect activities vary along altitudinal gradient; (3) evaluate how diversity and relative abundance of insect visitors influence reproductive output of *B. lycium* along altitudinal gradient. We discussed and suggested possible hypotheses that explain the pollination patterns observed as well as their implications in the context of the conservation of this substantially important medicinal plant.

## 2. Materials and Methods

### 2.1. Study area and flowering phenology

The present study was conducted at five different altitudes (800m, 1150m, 1500m 1850m, and 2200m; 33^º^32’-33^º^58’N and 74^º^32’-74^º^41’E) of Pir-Panjal mountain range in Jammu and Kashmir of north western Indian Himalaya from January, 2022 to August 2023. The Pir-Panjal mountain range splits up the Kashmir valley from the plains of the Jammu division (**Figure 1**). The largest mountain range in the lesser Himalayas is the Pir-Panjal range with varying average elevations from 1400 to 4100 m asl [2]. This altitudinal range offers the mountains and hillocks surrounding the Kashmir valley, and has enriched biodiversity above the elevation of 1700 m asl. The objective of this study embraces an diverse range of habitats which is reflected by a distinctive range of biodiversity distribution. The flowering and fruiting season of the plant extends from January–July. The flowers start appearing in January and continue up to June. The ripening of fruit begins in May and becomes fully matured up to early August at study sites along altitudinal gradients.

**Figure 1.**
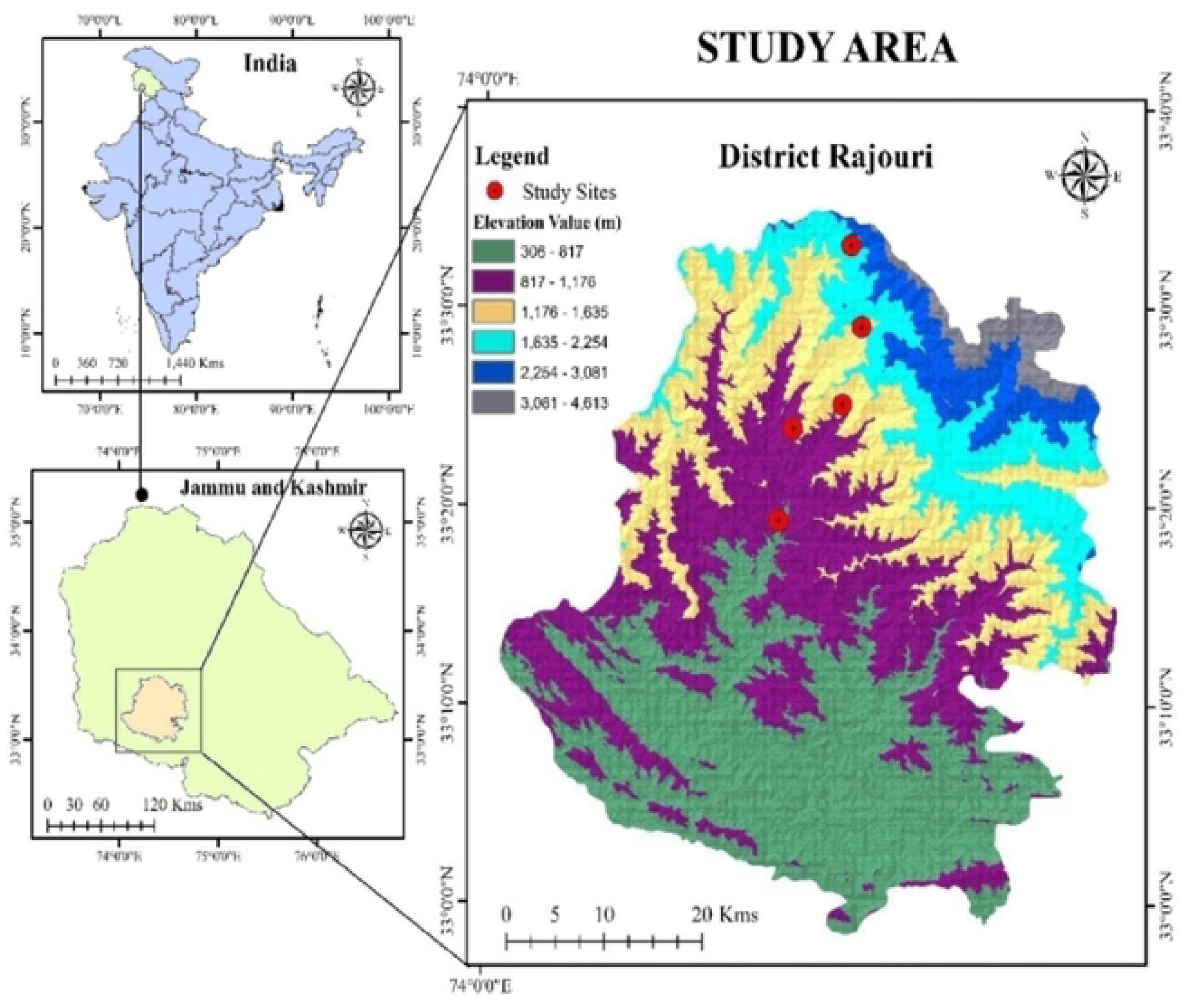
Map of the study area showing study sites along an altitudinal range.

### 2.2. Sampling and activities of insect pollinators

Documentation and monitoring of insect pollinators were conducted at the study sites between 08:00 h to 18:00 h from January to June when *B. lycium* was in full bloom for two consecutive flowering seasons during 2022-2023. In *B. lycium*, the efficiency of pollination by insects was monitored such as the category of insects visiting the plant, their foraging behavior, and the flower handling duration. To study the role that insects play in pollination, we used the insect net to catch notable flower visitor. Then we observed them under a stereozoom to examine the load of pollen on different parts of their body. For 30 plants of *B. lycium* in each site, three hours at various times of the day were devoted to observation and capturing the insect visitors to each plant. The specimens of the insect pollinators of *B. lycium* were identified at ITL (Insect Taxonomy Lab.), Department of Zoology, Baba Ghulam Shah Badshah University Rajouri. On each plant, we monitored 15-30 inflorescences, and total insect visitors in the observed inflorescences on 10 plants were considered in each site. During the peak flowering time,(mid-February up to early May) visiting insects were recorded at regular intervals of one hour throughout a 24-hour duration. To observe nocturnal foragers on *B. lycium*, a battery-operated torch was used. Due to the lack of nocturnal foragers on the plant, observations were limited from 0600 to 2000 hours (overall observation period =545 h).

The contribution of each pollinator to pollen transport was determined using various pollination indices:

#### 2.2.1. Foraging behaviour (FB)

It was calculated as the time spent by a specific pollinator per visit per inflorescence, counted by a stopwatch [32].

#### 2.2.2. Foraging speed (FS)

The number of flowers visited per minute by a pollinator is represented by FS. It was calculated as per [33]:

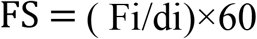

where ‘di’ is the time(s) shown by the stopwatch, and ‘Fi’ is the number of flowers visited during ‘di’.

#### 2.2.3. Index of visitation rate (IVR)

IVR is a relative measure of visitor rate that takes into consideration both activity rate and frequency of visits. It was calculated by following earlier work [32] as follows,

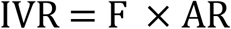

Where, ‘F’ represents the proportion of visiting-insect individuals to the total number of insects in the census, whereas ‘AR’ stands for activity rate, or the average number of flowers visited by a visiting-insect category in a minute.

#### 2.2.4. Insect visiting efficiency (IVE): IVE was calculated following [32]

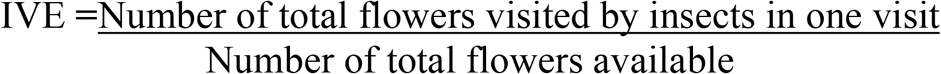

#### 2.2.5. Potential insect pollinator

To determine potential pollinators of *B. lycium*, the behaviour of insects was examined at two distinct behavioural decision levels.

##### (i) HRelative Abundance

The number of insect pollinators on *B. lycium* was observed on 15-30 inflorescences in each plant on a total of 30 plants in each site. The low or high abundance specified accessory, and major pollinators, respectively.

##### (ii) Sought floral resource

Insects were grouped on the basis of their purpose of visit on *B. lycium* like Collecting Pollen (CP), and Feeding on Pollen (FP), and Feeding on Nectar (FN).

### 2.3. Reproductive output

To calculate the reproductive output, the percentage of fruit and seed set by each plant was recorded. Percentage fruit set was determined by calculating the average number of flowers, and fruits per plant (Verma et al., 2021)[30]. Percentage fruit set was determined by using the formula:

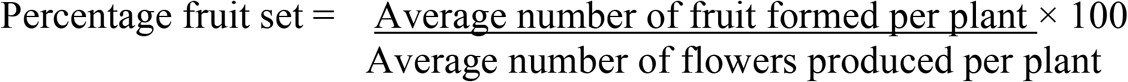

For the determination of percentage seed set, average number of ovules per ovary, and, average number of flower per plant was calculated [30]. Percentage seed set was calculated by using the formula:

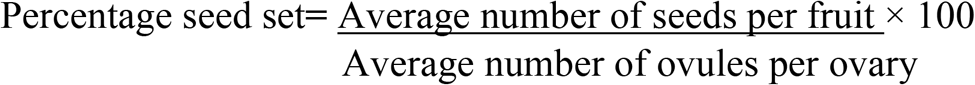

### 2.4. Data analysis

All the statistical analyses were carried out using R software version 4.0.2 (R Core Team 2020)[34]. Before data analysis, we evaluated whether the data on all studied pollination indices (foraging speed, foraging behavior, index of visitation rate, insect visiting efficiency, and relative abundance) met the requirement of homogeneity of variance and normality of distribution by using Levene’s test, and the Shapiro-Wilk normality test, respectively [35]. Since the assumptions of homogeneity of variance, and normality was not met for the studied pollination indices (*p*<0.001), we performed simple linear regression to study how insect activity and relative abundance as a function of altitude for all the order-wise data pooled together. Additionally, basic linear regression was used to examine the relationship between insect activity and pollen density. We checked the homogeneity and normality assumptions of the data using Levene’s test and the Shapiro-Wilk test before analyzing the variation in the insect pollination indices along the altitudinal gradient. In order to determine if there is a significant difference in the pollination indices along the altitudinal gradient, we used the unpaired two-sample T-test because the necessary assumptions have been met in the data (*p*>0.05).

## 3. Results

### 3.1. Plant species, floral arrangement and pollination

*B. lycium* is an evergreen, spiny, large shrub that grows at an altitudinal range between 800–2200 m asl. The leaves are coriaceous, slightly obovate-oblong or lanceolate having a few large, spiny teeth arranged alternately along the stem. The inflorescence is corymbose racemes containing 18-24 bright yellow, bracteate, hypogynous, actinomorphic, and hermaphrodite flowers that grow in axillary clusters. Over the calyx, two sporophylls are present. Six light yellow-colored sepals are present in two whorls, the outer three are smaller than the inner three. The corolla has six petals and is bright yellow. At the base of each petal, two nectaries are present. The androecium is composed of six antipetalous, adnate, and bithecous stamens. A single pistil is divided into the style, stigma, and ovary representing the gynoecium. The stigma has a central depression and is spherical. Style is short and empty. Pollination in *B. lycium*is carried out by insects as the plant possesses all the necessary entomophilous components, including brighter colour, scent, nectar, and pollen. Insect interaction is necessary for the pollen deposition on the stigmatic surface. The fruits are oval-shaped berries; acquire bright red or purple colour when fully ripened. Fruits contain 2-5 seeds which are yellow to pink in colour.

### 3.2. Species composition and activities of insect pollinators

In total, 1,934 individuals belonging to five orders, 17 families, and 39 species were found visiting *B. lycium* during the flowering period across altitudinal gradients. At site-1 (800 m), Diptera was the most dominant order with 8 species, followed by the Hymenoptera with four species, Lepidoptera, and Hemiptera each with 2 species, and Coleoptera with single species. At site-2 (1150 m), again Diptera was the most dominant order with 11 species, followed by Hymenoptera with 4 species, Hemiptera, Lepidoptera, and Coleoptera each having 2 species. At site-3 (1500 m), Diptera was the most dominant order again with 11 species, followed by Hymenoptera and Lepidoptera each with 3 species, and Hemiptera and Coleoptera each consisting of single species. However, at site-4 (1850 m), Lepidoptera was the most dominating order with 8 species followed by Diptera with 7 species, Hymenoptera with 5 species and, Coleoptera with 2 species. At site-5 (2200 m), Lepidoptera and Hymenoptera both consisting of 7 species followed by Hymenoptera with 3 species and Coleoptera with 2 species (Table 1 and Figure 2 and 3).

**Table 1.** Showing insect pollinator species belonging to different orders, their foraging activity and altitudinal range (FN= Feeding on Nectar, CP= Collecting Pollen, and FP=Feeding on Pollen).

| Species | Order | Family | Foraging activity | Altitude (m) |
| --- | --- | --- | --- | --- |
| *Allograpta* sp. | Hymenoptera | Apidae | FN | 800 |
| *Apis* sp. | Hymenoptera | Apidae | FN, CP, FP | 800-2200 |
| *Bombus trifasciatus* | Hymenoptera | Apidae | FN | 800-1150 |
| *Formica fusca* | Hymenoptera | Formicidae | FP | 800-1150 |
| *Bombus* sp. | Hymenoptera | Apidae | FN, CP | 1850-2200 |
| *Camponotuspennsylvanicus* | Hymenoptera | Formicidae | FN, CP, FN | 1500-2200 |
| *Bombus haemorrhoidalis* | Hymenoptera | Apidae | FN, CP | 1850 |
| *Apis cerena* | Hymenoptera | Apidae | FN, CP, FP | 1500-1850 |
| *Coelioxys* sp. | Hymenoptera | Megachilidae | FN, CP | 800 |
| *Episyrphusbalteatus* | Diptera | Syrphidae | FP | 800-2200 |
| *Eristalistenax* | Diptera | Syrphidae | FP | 800-2200 |
| *Musca domestica* | Diptera | Muscidae | FP | 800-2200 |
| *Calliphora vomitoria* | Diptera | Calliphoridae | FN, CP, FP | 800-1150 |
| *Chrysotoxumbaphyrum* | Diptera | Syrphidae | FN | 800-1150 |
| *Eupeodesluniger* | Diptera | Syrphidae | FP | 800-1500 |
| *Sarchophaga* sp. | Diptera | Sarchophagidae | FP | 800-1500 |
| *Plecia* sp. | Diptera | Bibionidae | FP | 800-1150 |
| *Rhingi*asp. | Diptera | Syrphidae | FN, FP | 1150-1500 |
| *Calliphora*asp | Diptera | Calliphoridae | FN, CP, FP | 1150 |
| *Eristaliscereal*is | Diptera | Syrphidae | FP | 1150-2200 |
| *Episyrphysviridaureus* | Diptera | Syrphidae | FP | 1500-2200 |
| *Eristalinustaeniops* | Diptera | Syrphidae | FP | 1550 |
| *Lucilia* sp. | Diptera | Calliphoridae | FN, CP, FN | 1500-2200 |
| *Syritta* sp. | Diptera | Syrphidae | FP | 1500-2200 |
| *Dyscedrus*sp. | Hemiptera | Coreidae | FP | 800-1150 |
| *Aphidoidea* | Hemiptera | Aphididae | FP | 800-1500 |
| *Praezygaenacaschmirensis* | Lepidoptera | Zygaenidae | CP, FP | 800-1500 |
| *Celastrinaargiolus* | Lepidoptera | Lycaenidae | FN | 800-2200 |
| *Dodona durga* | Lepidoptera | Riodinidae | FN | 1500-2200 |
| *Heliophorusmoorei* | Lepidoptera | Lycaenidae | FN | 1850-2200 |
| *Celastrina* sp. | Lepidoptera | Lycaenidae | FN | 1850 |
| *Heliophorus sena* | Lepidoptera | Lycaenidae | FN | 1850-2200 |
| *Heliophorus*sp. | Lepidoptera | Lycaenidae | FN | 1850 |
| *Cyrestisthyodamas* | Lepidoptera | Nymphalidae | FN | 1850-2200 |
| *Aglaiscaschmirensis* | Lepidoptera | Nymphalidae | FN | 1850-2200 |
| *Pieris* sp. | Lepidoptera | Pieridae | FN | 2200 |
| *Coccinellaundecimpunctata* | Coleoptera | Coccinellidae | CP | 800-2200 |
| *Altica* sp. | Coleoptera | Chrysomelidae | FN, FP | 1150-1500 |
| *Coccinella septempunctata* | Coleoptera | Coccinellidae | CP | 1850-2200 |

**Figure 2.**
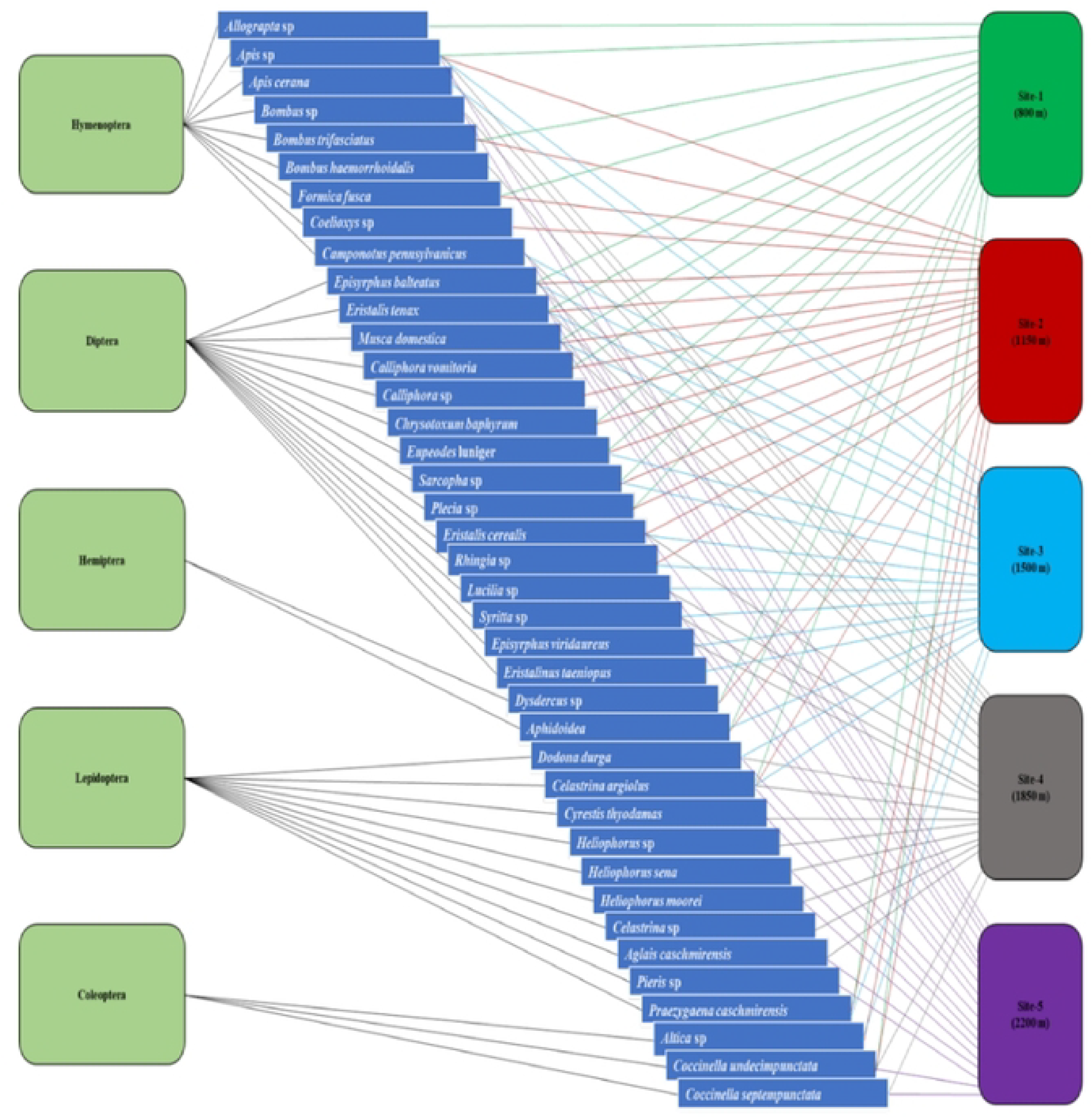
Interaction of insect pollinators of *Berberis lycium* across study sites along altitudinal gradient.

**Figure 3.**
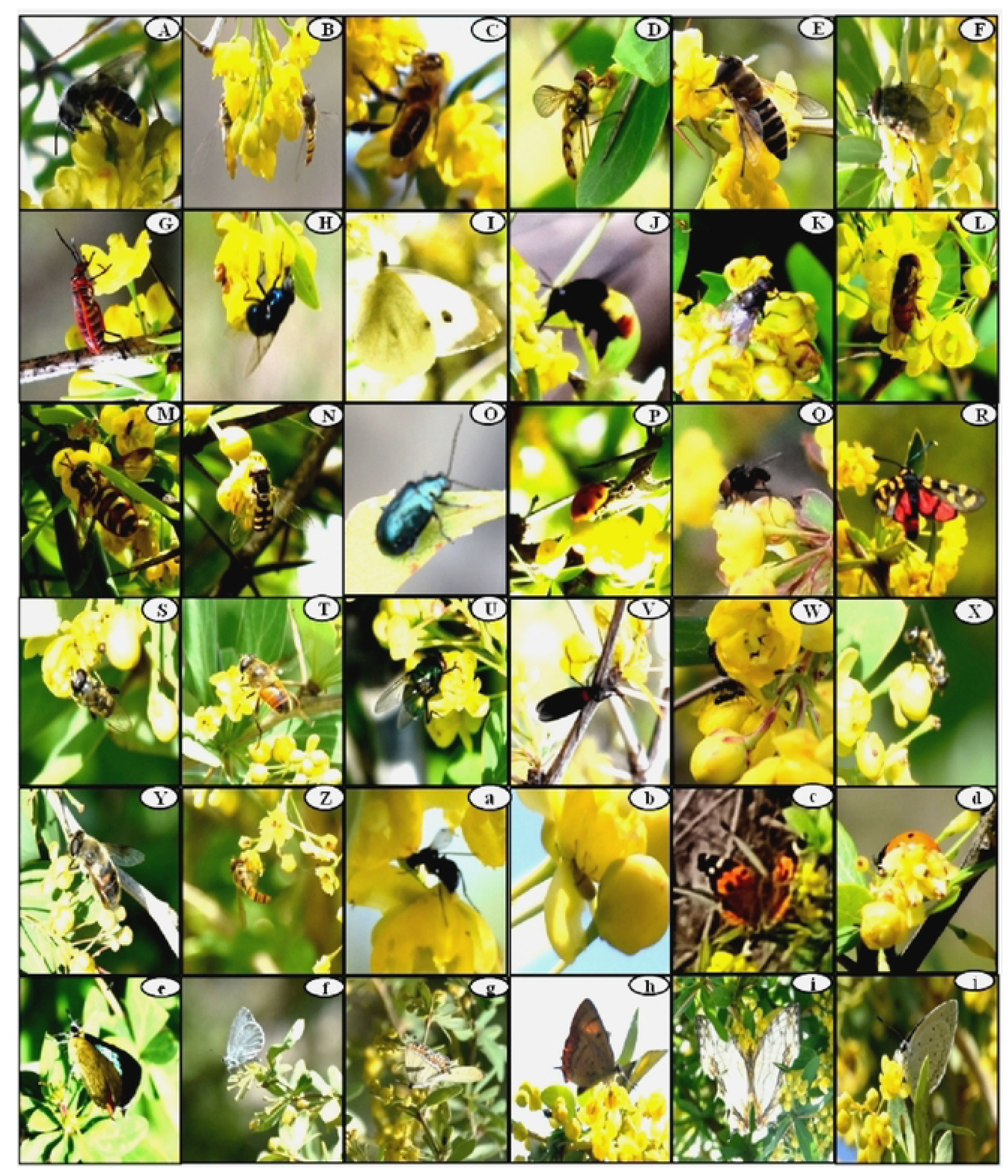
Representing specimens of **v**arious insect pollinators on *Berberis lycium*flowers at different altitudinal range of the study area: A) *Apis* sp., B) *Episyrphusbalteatus*, C) *Apis cerena*, D) *Allograpta* sp., E) *Eristaliscerealis*, F) *Musca domestica*, G) *Dysdercus*sp, H) *Calliphora vomitoria*, I) *Pieris* sp., J) *Bombus trifasciatus*, K) *Calliphora* sp., L)*Rhingia* sp., M) *Chrysotoxumbaphyrum*N) *Eupeodesluniger*, O)*Altica* sp., P)*Coccinellaundecimpunctata*, Q) *Sarcopha*sp., R)*Praezygaenacaschmirensis*, S) *Syritta* sp., T) *Eristalistenax*, U) *Lucilia* sp., V) *Plecia* sp., W) *Formica fusca*, X) *Eristalis* sp., Y) *Eristalinustaeniops*, Z) *Episyrphusviridaureus*, a) *Camponotuspennyslyvanicus*, b) *Aphidoidea*, c)*Aglaiscaschmeriensis*, d) *Coccinellaseptempunctata*, e) *Heliophorusmoorei*, f) *Celastrina*sp, g*) Heliophorus sena*, h*) Heliophorus* sp., i) *Cyrestisthyodamas*, j)*Celastrinaargiolus*.

### 3.3. Insect activities

The results of the linear regression showed a statistically non-significant increase (*p*>0.05) in terms of foraging behaviour of all the studied insect pollinator orders with the highest increase in Diptera and Hymenoptera with an increase in altitude respectively (Figure 4A).

**Figure 4.**
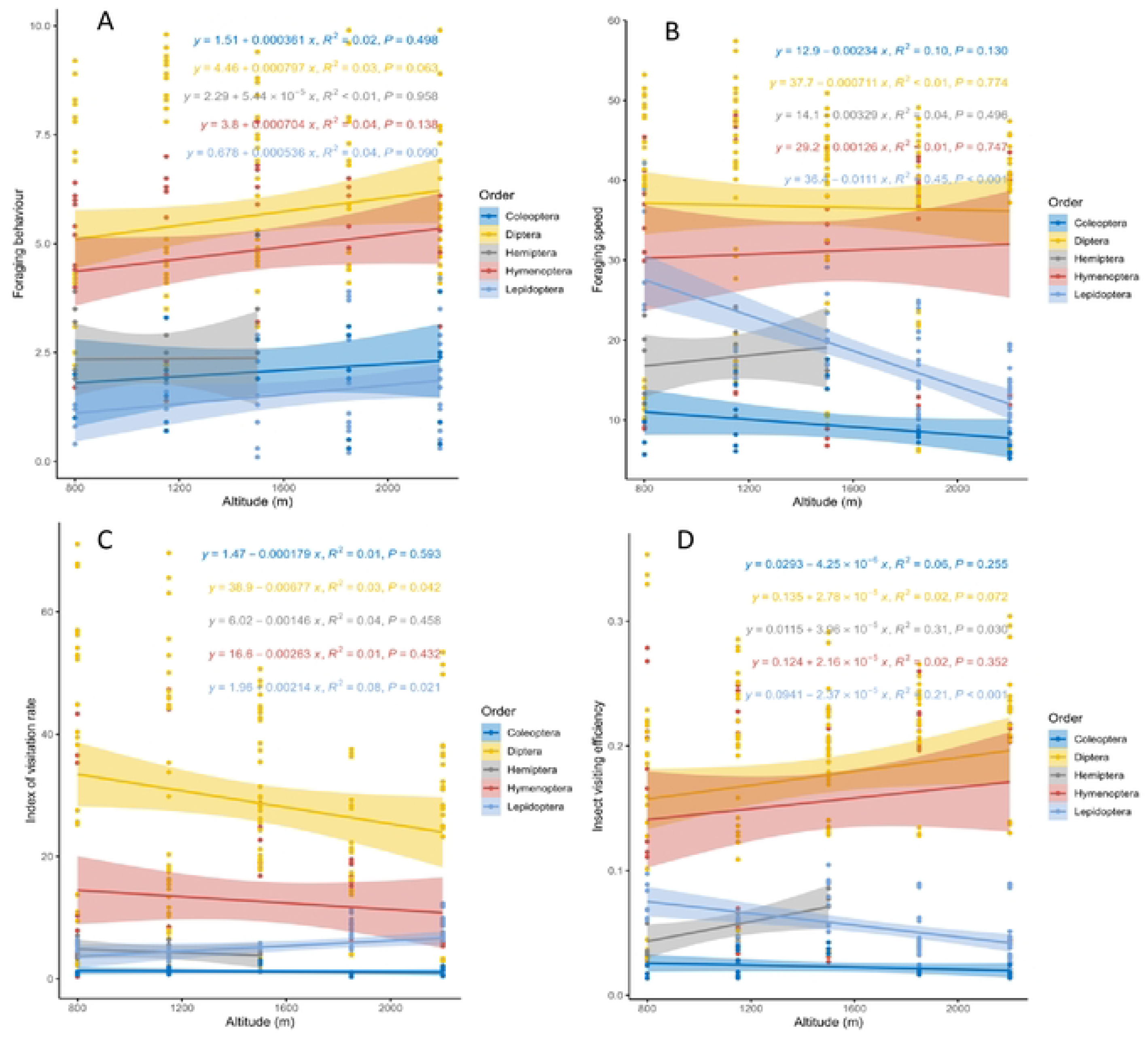
Activities recorded of different pollinators’ orders at different altitudes, A. Foraging behaviour, B. Foraging speed, C. Index of visitation rate, D. Index of visiting efficiency. The best-fitting regression line is represented by the coloured solid line, while the 95% confidence interval of regression line is shown by the shaded areas.

We found a statistically significant decrease (*p*<0.0001) in the foraging speed of the order Lepidoptera only (y=36.4-0.0111x, R^2^ =0.05). However, a statistically non-significant increase (*p*>0.05) in Hymenoptera and Hemiptera and a non-significant decrease (*p*>0.05) in Coleoptera and Diptera have been observed in the studied insect orders with an increase in altitude (Figure 4B).

Further, with an increase in altitude, a statistically significant decrease (*p*<0.05) in insect visitation rate has been observed for the order Diptera (y=38.9-0.00677x, R^2^=0.03), whereas, a statistically significant increase (*p*<0.05) for insect visitation rate has been observed for the order Lepidoptera (y=1.96+0.00214x, R^2^=0.08). However, a statistically non-significant decrease (*p*>0.05) in terms of IVR has been observed for Coleoptera, Hemiptera, and Hymenoptera insect orders (Figure 4C).

Similarly, a statistically significant increase (*p*<0.05) in insect visiting efficiency was observed for the order Hemiptera (y=0.0155+3.96×10^−5^x, R^2^=0.31) with an increase in altitude. Whereas, a statistically significant decrease (*p*<0.001) has been observed for the order Lepidoptera (y=0.0941-2.37×10^−5^x, R^2^=0.21). However, a statistically non-significant decrease (*p*>0.05) was observed for Coleoptera, and the non-significant increase was observed for Diptera and Hemiptera insect orders with an increase in altitude (Figure 4D).

We found a statistically significant decrease (*p*<0.005) for foraging speed (y=36.6-0.00566x, R^2^=0.03) and index of visitation rate (y=26.4-0.0061x, R^2^=0.03) in overall insect species with an increase in attitude. However, a statistically non-significant decrease (*p*<0.05) was observed in the foraging behaviour and insect visiting efficiency with an increase in altitude (Figure 5).

**Figure 5.**
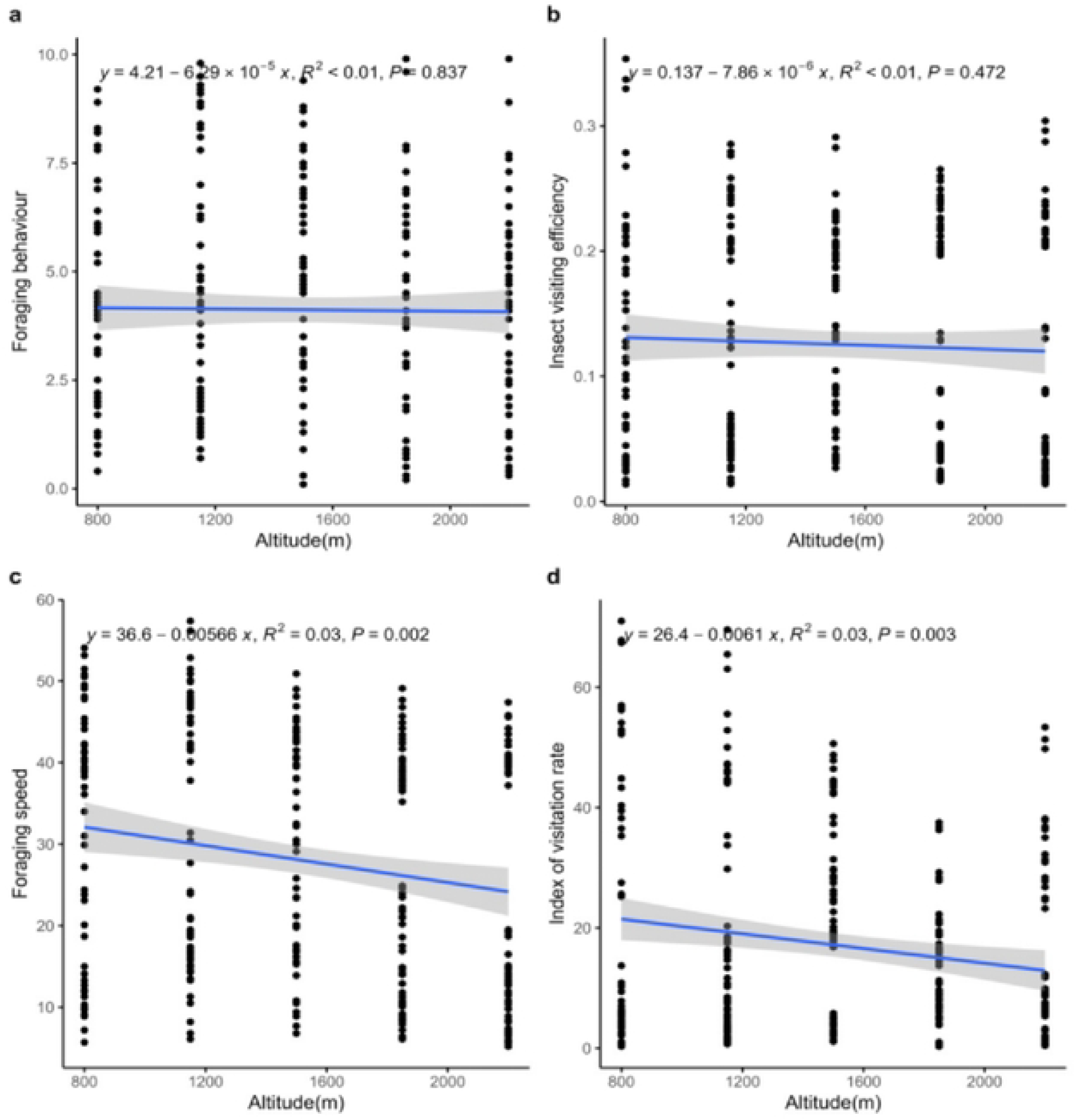
Activities recorded of combined pollinators’ orders at different altitudes, A. Foraging behaviour, B. Foraging speed, C. Index of visitation rate, D. Index of visiting efficiency. The best-fitting regression line is represented by the blue solid line, while the 95% confidence interval of regression line is shown by the shaded areas.

Then, we found a statistically significant increase (*p*<0.05) in relative abundance in the case of Lepidoptera (y=0.929+0.00264x, R^2^=0.31). However, a statistically non-significant decrease (*p*<0.05) was observed in Diptera and Hemiptera and a non-significant increase in Coleoptera and Hymenoptera was observed with an increase in altitude (Figure 6).

**Figure 6.**
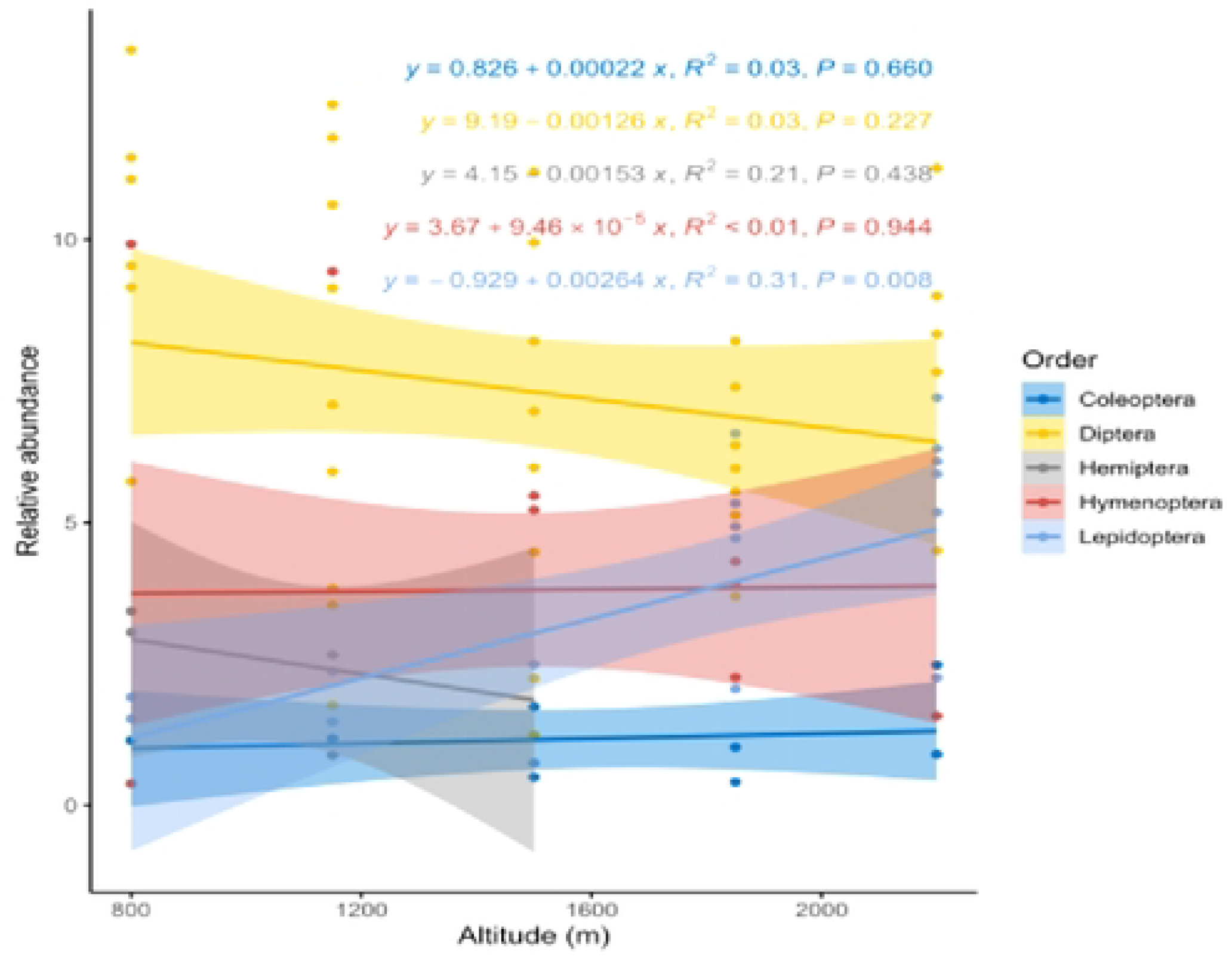
Relative abundance of different pollinators’ orders at different altitudes. The best-fitting regression line is represented by the coloured solid line, while the 95% confidence interval of regression line is shown by the shaded areas.

The average percentage fruit set and percentage set decreased with increasing elevation (Figure 7), and the decrease was statistically non-significant (*p*<0.05) in the case of the percentage fruit set (y=71-0.0154 x, R^2^=0.43) and, percentage seed set (80.9-0.0104 x, R^2^=0.42) with an increase in altitude.

**Figure 7.**
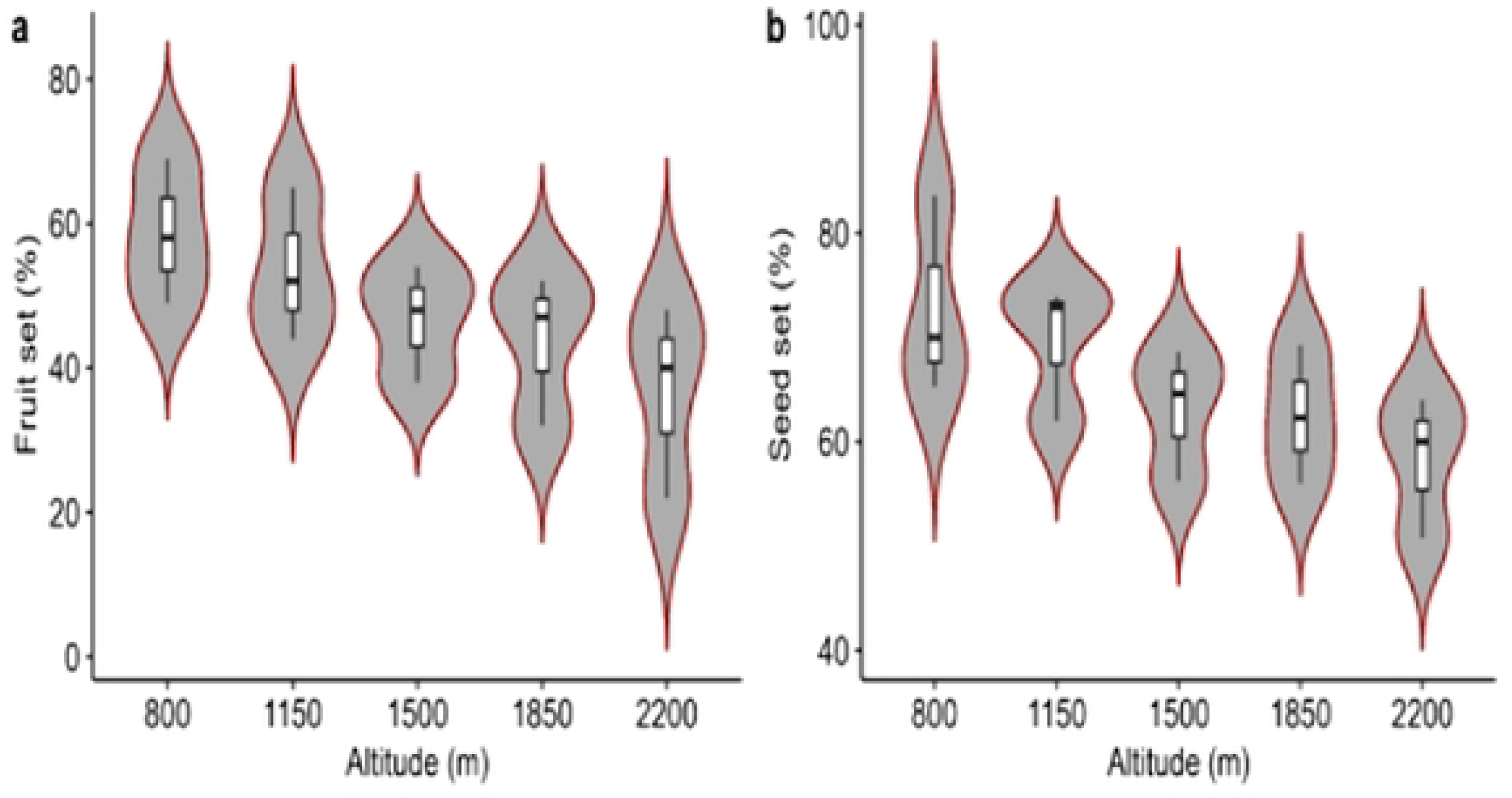
Change in reproductive output along altitudinal gradient. a) fruit set (PFS; %), b) seed set (PSS; %), The shape in violin plots at 95% confidence, indicates the data of reproductive output. The violin plots are escorted by whisker boxplots, in which the average reproductive output for each parameter is shown by black lines.

## 4. Discussions

In this study, we documented various types of insect pollinators along altitudinal gradients commonly found foraging on flowers of *B. lycium*. In total 1,934 individuals of insect pollinators belong to 39 species under five orders and 17 families were observed visiting the plant species during the flowering period across altitudinal gradient. Pooled insect order data in each altitude were used to assess differences in insect activities and abundance across elevation zones. We observed significant variations in the insect pollination indices along five altitudinal gradients. Our results also indicated differences in the relative abundance of insect pollinators and reproductive output along the altitudinal gradients. However, the repetition of collections in various sampling days in the individual sites offers a broad assessment of species flowering phenology and insect visitors along altitudinal gradients with greater precision.

This is the first study to document potential insect pollinators of *B. lycium* along the altitudinal gradient in the Pir-Panjal region of Indian western Himalaya. We found that *B. lycium* has a more diverse assemblage of pollinators as previously reported [25, 30] due to a wide range of altitudinal gradients. In the case of foraging behavior (FB), the *Apis*species is the most common forager reported in all the studied altitudes (Table 1). However, *Musca domestica, Calliphora vomitoria, Sarchophaga*sp. and *Lucilia* sp. were reported to have the highest foraging behaviour among Hymenoptera. The foraging behaviour of all the studied insect pollinator orders shows a statistically non-significant increase with the highest increase in Diptera and Hymenoptera with an increase in altitude respectively (Figure 4A). In case of foraging speed (FS), *Calliphora vomitoria, Bombus trifasciatus, Sarchophaga*sp. and,*Eupeodesluniger*have the highest foraging speed. However, among all the studied orders a statistically significant decrease in foraging speed was observed in Lepidoptera only (Figure 4B). In the case of Insect visiting efficiency (IVE), *Bombus trifasciatus, Calliphora vomitoria*,and *Episyrphusbalteatus*showthe highest activity. When data were polled together, only Diptera showed a significant decrease and Lepidoptera shows significant increase in the insect visiting efficiency (IVE) with the increase in altitude respectively (Figure 4C). Further, in index of visitation rate (IVR), *Calliphora vomitoria, Eupeodesluniger, Episyrphusbalteatus*, and *Eristalistenax*shows highest activity. When data polled together, only Lepidoptera shows significant decrease in the (IVR) with the increase in the altitude (Figure 4D). While considering all the four parameters (FB, FS, IVE and, IVR) and data pooled together, the activity rate shows a decrease with an increase in altitude. However, only foraging speed (FS) and index of visitation rate (IVR) show statistically significant results (Figure 5). Lastly, in the case of relative abundance of insect pollinators, Lepidoptera shows a statistically significant increase with the increase in altitude (Figure 6). In the final important observation, we observed a statistically significant decrease in fruit and seed set along the altitudinal gradient (Figure 7).

Observations on major pollinator groups (Hymenoptera, Diptera, Coleoptera, Lepidoptera) may serve as efficient indicators of changes in micro-climate that directly affect their foraging and nesting place [36]. The majority of Pollinators belonging to Lepidoptera show a distinct response to altitude in which they showed their presence in the elevation range between 1500-2200m with few exceptions which signifies its importance in climate change studies [37]. The members of Hemiptera were confined to lower altitudes only; hence they were not considered as potential pollinators of *B. lycium*. The members of Hymenoptera and Diptera showed their presence in a wide range of altitudes. At the order level and, at the base altitude of 800 m, flies (Diptera) predominate (with few exceptions) as potential visitors of *B. lycium* (Table 1). Above 800 m the most prominent flower visiting orders altered among sites with no consistent altitudinal pattern except Lepidoptera. Flies (Diptera) therefore represented comparatively diverse and abundant potential visitors that provide pollination services to *Berberis* to a greater extent [25, 30, 31, 38], and decrease in their abundance and activity along altitudinal gradient can be linked to a decrease in the reproductive output of this essentially important medicinal plant.

Altitudinal variations in insect taxonomic diversity can be observed in varied spatial scales [39]. On a finer scale, the number of insect pollinators associated with a specific plant species is reported to decrease with increasing altitude [6, 8, 40, 41] reported a decline in the insect fauna of *Betula pubescens* (Heteroptera, Coleoptera, and Homoptera species) from 0 to 900 m in Norway. Another study conducted on *Campanula rotundifolia* showed comparatively lower pollinator diversity and visitation rate of insect visitors at higher elevations as highly efficient bumblebees replace less efficient solitary bees in higher elevations [42]. These previous observations are in parallel with our findings as all observed activities of insect visitors show a decrease with increasing altitude. However, only foraging speed and index of visitation rate show statistically significant results (Figure5). Our results indicated that the foraging behaviour of visiting insects of all studied orders increases with increasing altitude (Figure 4A). This variation in foraging behaviour also reflects host plant abundance and richness [14] and also due to the decline in interacting partners at high elevations (Adedoja et al., 2018)[43]. Butterflies and moths are diverse, the most studied insects and highly sensitive to climate, therefore become ideal for climate change research [37]. This coincides with our findings because only Lepidoptera shows a significant decrease in foraging speed with an increase in altitude (Figure 4B). Our work (supplementary table S1) coincides with the previous findings [44], which observed a positive correlation between the diversity of Bumblebees with increasing altitude and a negative correlation between the diversity and visitation rate of non-bumblebees with increasing altitude. In the case of insect visiting efficiency, only Lepidoptera and Hemiptera show significant decrease and increase in activity respectively (Figure 4D). At higher altitudes of temperate regions, anthophilous flies are often the sole pollinators [45-47] but at lower altitudes presence of Hymenoptera increases. Hemiptera, in our study, plays a minor role and tends to decrease with increasing altitude.

A great diversity of insects are specialized for feeding on host plant reproductive and floral structures [32, 48, 49], but many plant species experience a declined reproductive output with increasing elevation [6, 27, 50, 51]. The fitness of plant-pollinator interactions is essential for plant reproductive assurance and the evolution of floral traits and breeding systems [35]. Breeding systems usually vary within or among different plant lineages that include complete as well as intermediate outcrossers, and selfers [52, 53] and lineages undergo repeated selfing may often suffer the loss of fitness due to inbreeding depression. Therefore, insect pollination has been proven to be a key factor for both the nutritional and, physical quality [54]. Although biotic interactions play a significant role in limiting the expansion of many plant species, pollination syndrome also plays a significant evolutionary function in maintaining ecological range boundaries [55]. Insect pollinator decline is likely to affect plant reproductive traits that grow in vulnerable habitats. For instance, early flowering, and self-incompatible plants are more reliant on pollinators than mid-flowering, and self-compatible plants, therefore pollinator decline is expected to have a significant impact on these types of plants. The high reliance of *B. lycium* on pollinators as observed in our study leads us to speculate that the high dependence on pollinators for the production of seed observed in our study suggests that if plants cannot adjust their reproductive strategies to reduce this reliance amidst pollinator declines, their seed production will be drastically affected by climate change [56].

## 5. Conclusion

For conservation biologists looking at how plant-pollinator interactions may be affected, our findings could be helpful as global climate change is predicted to alter spatio-temporal distribution and interactions of anthophilous insects. Butterflies and moths (Lepidoptera) appear to be the most sensitive pollinators and show significant variations in their activity along altitudinal gradients. We documented the diversity of anthophilous insects associated with the pollination of *B. lycium* along altitudinal gradients. As a global trend, our study also observed the decrease in the overall abundance and activities of insect pollinators of *B. lycium* with an increase in altitude have a negative correlation with its reproductive output. The high dependence of *B. lycium* on pollinators, as observed in our study, suggests that if plants cannot adjust their reproductive strategies to reduce this reliance amid pollinator declines, their seed production will be significantly impacted by climate change.

## Supporting information

**Supplementary table S1**. Showing Foraging Behaviour, Foraging Speed, Insect Visiting Efficiency, Index of Visitation Rate, Density and Relative abundance of various individual insect species and orders at different altitudes.

## Author Contributions

**Conceptualization:** Nahila Anjum, Susheel Verma, Mohd Hanief

**Data curation:** Nahila Anjum

**Formal analysis:** Nahila Anjum, Sajid Khan

**Funding acquisition:** Nahila Anjum

**Investigation:** Susheel Verma, Kailash S Gaira, Balwant Rawat, Nakul Chettri, Mohd Hanief

**Methodology:** Nahila Anjum, Sajid Khan

**Project administration:** Susheel Verma, Nakul Chettri, Mohd Hanief

**Resources:** Nahila Anjum, Susheel Verma, Mohd Hanief

**Software:** Nahila Anjum, Sajid Khan

**Supervision:** Nakul Chettri, Mohd Hanief

**Validation:** Susheel Verma, Kailash S Gaira, Balwant Rawat, Nakul Chettri, Mohd Hanief

**Visualization:** Balwant Rawat

**Writing – original draft:** Nahila Anjum, Sajid Khan

**Writing – review & editing:** Nahila Anjum, Sajid Khan, Susheel Verma, Kailash S Gaira, Balwant Rawat, Nakul Chettri, Mohd Hanief

## Acknowledgments

The authors thank Ms. Taslima Sheikh and Dr. Sajad H. Parey, Department of Zoology, BGSB University, Rajouri for insect identification. Besides, first author (Ms. Nahila Anjum) is thankful to CSIR-HRDG, New Delhi for providing Junior and Senior Research Fellowship (File no. 09/1172(0005)/2019-EMR-I).

